## Supplementary Figures for "Structure and function of bacterial YeeE-YeeD complex in thiosulfate uptake pathway"

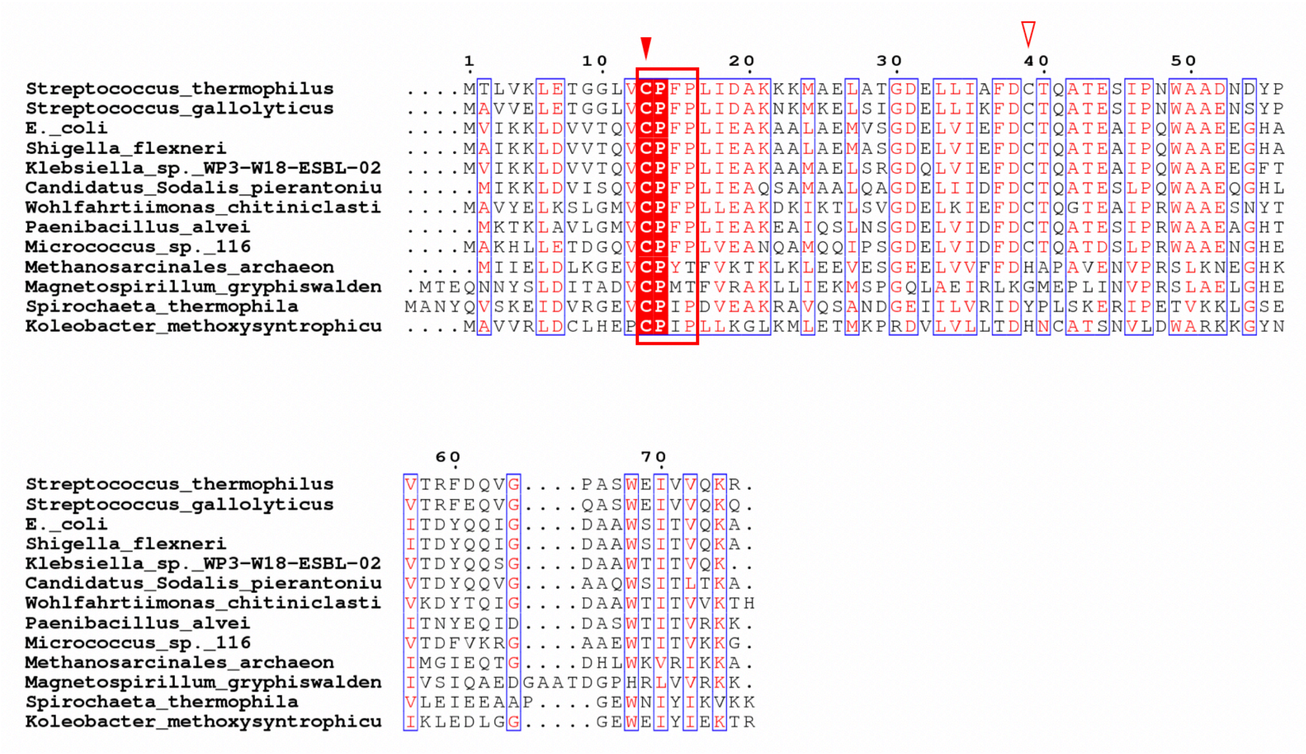


**Fig. S1. Sequence alignment of YeeDs from thirteen species.**

The UniProt IDs of YeeD sequences used are as follows: *E. coli* YeeD, P33014; *Spirochaeta_thermophila* YeeD, G0GAP7; *Candidatus Sodalis pierantonius* YeeD, W0HL40; *Klebsiella_sp._WP3-W18-ESBL-02* YeeD, A0A7I6Q8F8; *Methanosarcinales archaeon* YeeD, A0A822J3Z6, *Streptococcus thermophilus* YeeD, A0A8D6XUG1; *Koleobacter methoxysyntrophicus* YeeD, A0A8A0RNL6; *Wohlfahrtiimonas chitiniclastica* YeeD, L8Y0N6; *Paenibacillus alvei* YeeD, A0A383RJV0; *Shigella flexneri* YeeD, A0A384L8W9; *Streptococcus gallolyticus* YeeD, A0A380K504; *Micrococcus_sp._116* YeeD, A0A653IT90; *Magnetospirillum gryphiswaldense* YeeD, V6F4H7. The red rectangle indicates the CPxP motif. The first and second cysteine residues (solid and open red arrowheads) are completely and not completely conserved among species, respectively. The figure was generated using ESPript 3.0 (*45*).

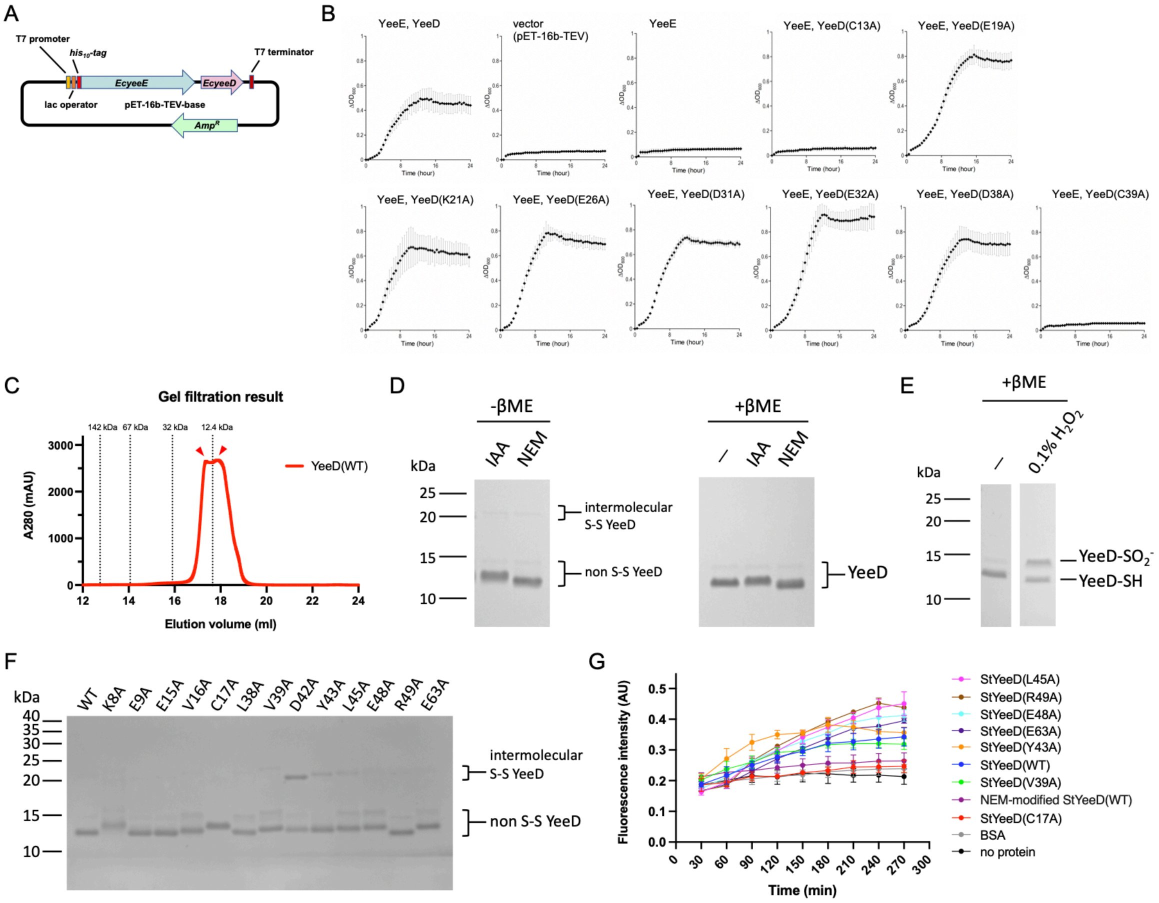


**Fig. S2. Data related to functional analysis of YeeD.**
(**A**) Details of plasmid (pAZ061), used as the positive control for the growth complementation assay. Tandemly located *EcyeeE* and *EcyeeD* are regulated by the same promoter. A His$_{10}$-tag is attached to the N-terminal side of *Ec*YeeE. Based on pAZ061, a deletion and several point mutations on *EcyeeD* were introduced. (**B**) Original data from the growth complementation assay in Fig. 2C. Error bars represent the SD of three measurements. (**C**) Gel filtration profile of purified *St*YeeD(WT), which eluted with two peaks (red arrowheads). The eluted positions of standard proteins and their molecular masses are shown. (**D**) Non-reducing (-βME) and reducing (+βME) SDS-PAGE of *St*YeeD after iodoacetamide (IAA)- or N-Ethylmaleimide (NEM) treatment. Only minor fractions show intermolecular disulfide bond formation between *St*YeeDs(WT). **(E)** Irreversible oxidation of *St*YeeD by hydrogen peroxide. Before reducing SDS-PAGE, the hydrogen peroxide treatment was performed. **(F)** Non-reducing SDS-PAGE profile of *St*YeeD mutants after NEM treatment. **(G)** Original data for the enzymatic activity of *St*YeeD detected by HSip-1 in Fig. 2H. Error bars show the SD from three measurements.

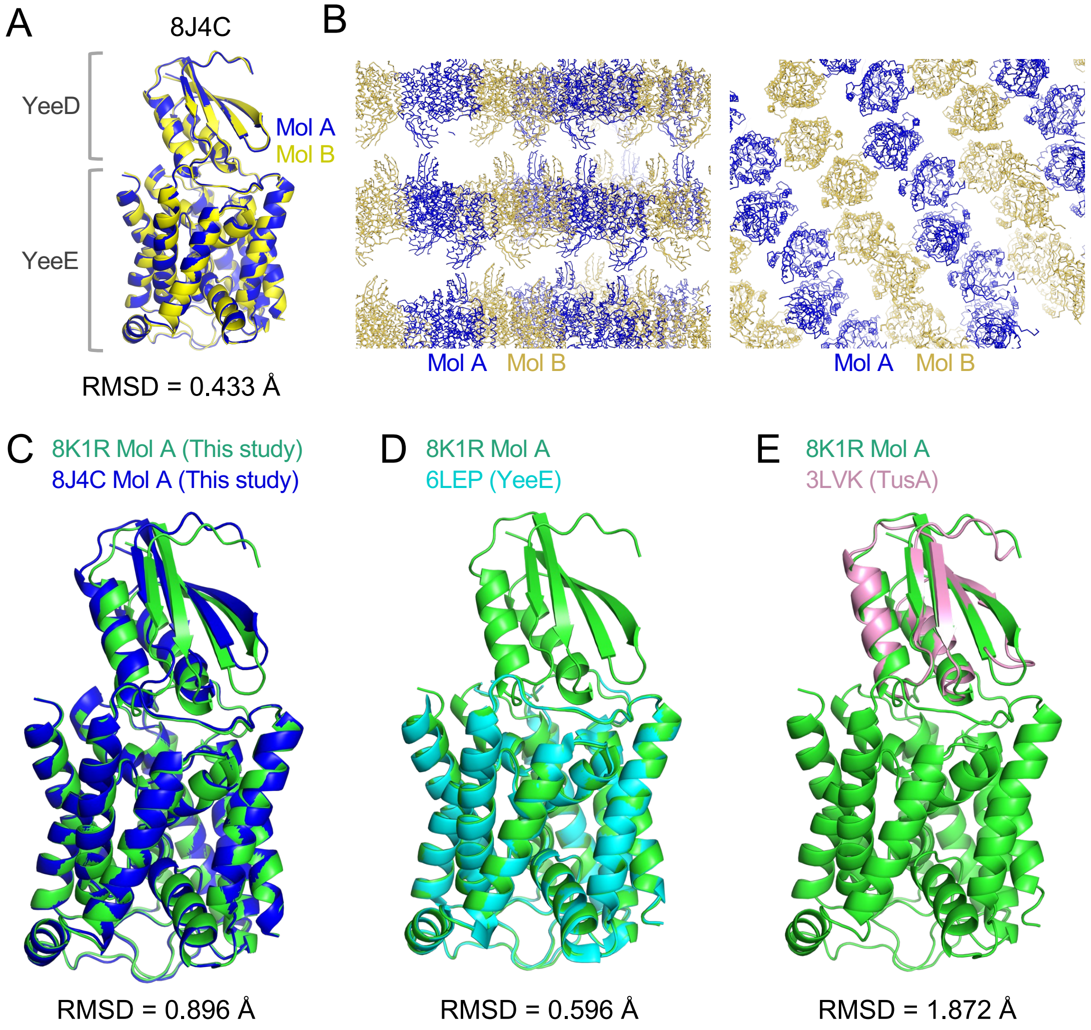


**Fig. S3. Structural comparisons of *St*YeeE-YeeD complex.**

(**A**) A comparison of the crystal structures Mol A and Mol B in the asymmetric unit of 8J4C. (**B**) Crystal packing of 8J4C viewed from two different directions. (**C**) Comparison of the crystal structures of Mol A of 8K1R and MolA of 8J4C. (**D** and **E**) Comparisons of the crystal structures of *St*YeeE-YeeD complex (MolA) and *St*YeeE (PDB ID 6LEP) (D) or *E. coli* TusA (PDB ID 3LVK) (E).

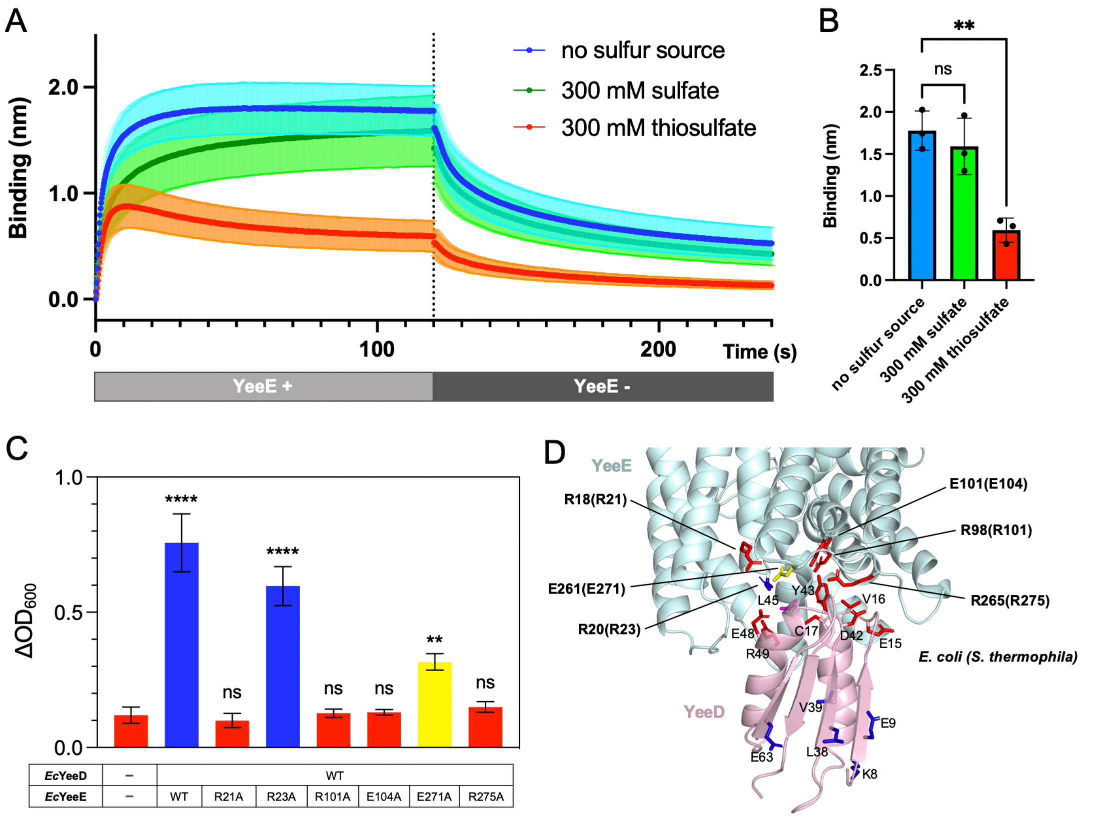


**Fig. S4. Data related to interactions between YeeE and YeeD.**
(**A**) Real-time detection by the BLI method of association and dissociation of *St*YeeE with/from solidified *St*YeeD in buffer without a sulfur source, with 300 mM $Na_2SO_4$, or with 300 mM $Na_2S_2O_3$. Each line shows the mean value of three measurements with the SD. The dashed line indicates the 120-s point. (**B**) Comparison of mean values of *St*YeeE binding to solidified *St*YeeD at 120 s. Error bars represent the SD of three measurements. Statistical significance compared with no sulfur source was determined using one-way analysis of variance (ANOVA) followed by Dunnett's multiple comparisons tests (**, $p < 0.01$; ns, not significant). (**C**) Growth complementation assay of Δ*cysPUWA*Δ*yeeE* (DE3) cells, depending on *Ec*YeeE and *Ec*YeeD expressed from plasmids. Mean values of increased $OD_{600}$ ($\Delta OD_{600}$) after 24 h are shown. Error bars represent the SD from three measurements. Statistical significance compared with the Δ*cysPUWA* Δ*yeeE* (DE3) cells possessing an empty vector was determined using one-way ANOVA followed by Dunnett's multiple comparisons tests (****, $p < 0.0001$; **, $p < 0.01$; ns, not significant). (**D**) Mutation site mapping on the crystal structure of *St*YeeE-YeeD complex (PDB ID 8J4C). The side chains of amino acid residues of *St*YeeE corresponding to those of *Ec*YeeE mutated in (C) are shown as stick models with the same colors as in (C). The amino acid residues of *St*YeeD shown in Fig. 4C are also indicated.

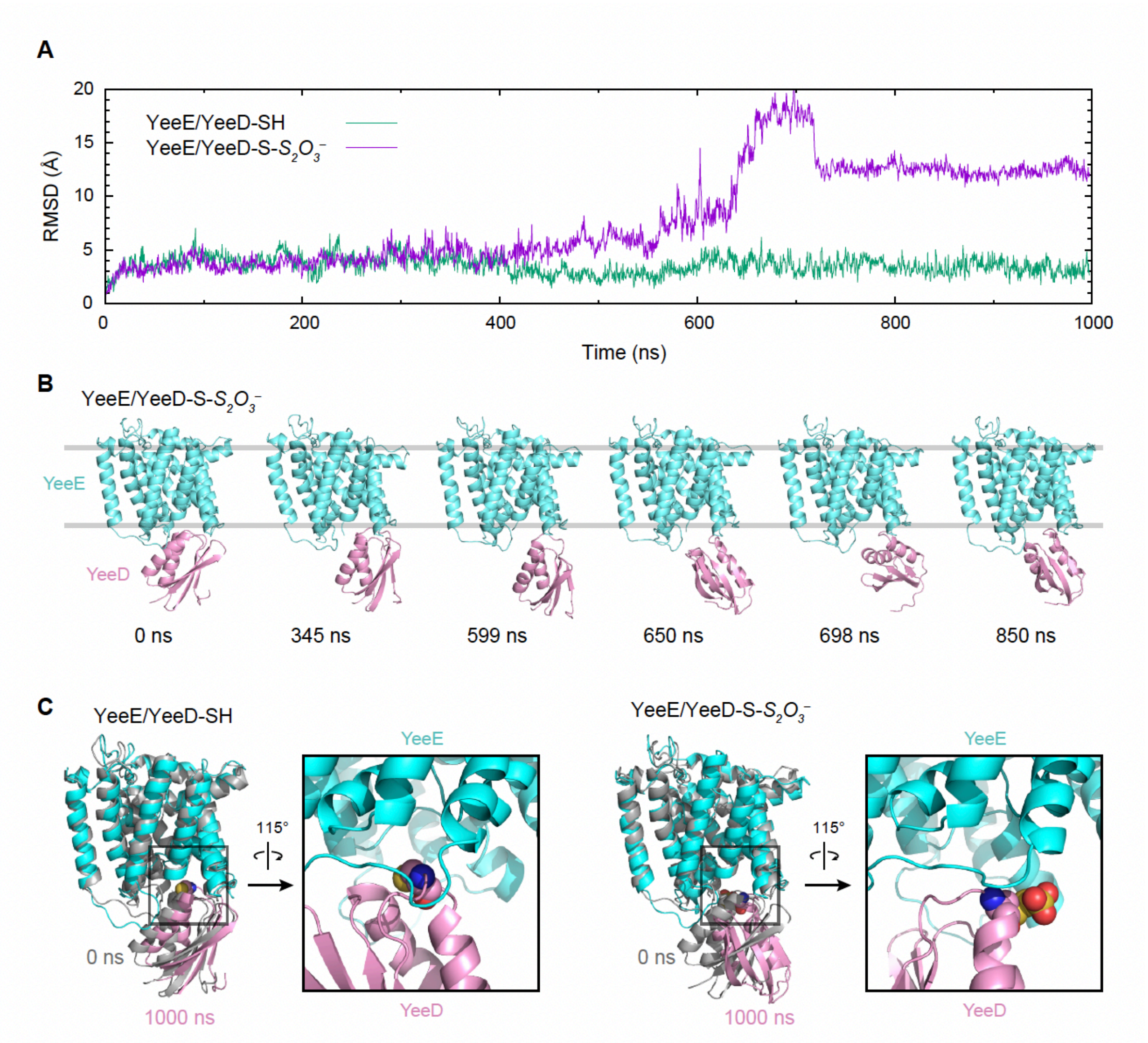


**Fig. S5. All-atom MD simulations of *St*YeeE/YeeD-SH and *St*YeeE/YeeD-S-$S_2O_3^-$**

(**A**) Time courses of the Cα-RMSD calculated for *St*YeeD regions relative to the initial structures. For the RMSD calculations, each *St*YeeE region was superimposed. (**B**) Snapshots in the MD simulation of *St*YeeE/YeeD-S-$S_2O_3^-$. (**C**) Comparison between the initial structure (0 ns) and final snapshot (1000 ns) for *St*YeeE/YeeD-SH (left) and *St*YeeE/YeeD-S-$S_2O_3^-$ (right). The square boxes display magnified views of the region around C17 (spheres).
